## Supplemental figures for "SUMOylation of Bonus, the *Drosophila* homolog of Transcription Intermediary Factor 1, safeguards germline identity by recruiting repressive chromatin complexes to silence tissue-specific genes"

### Extended Data Fig.1

**A**

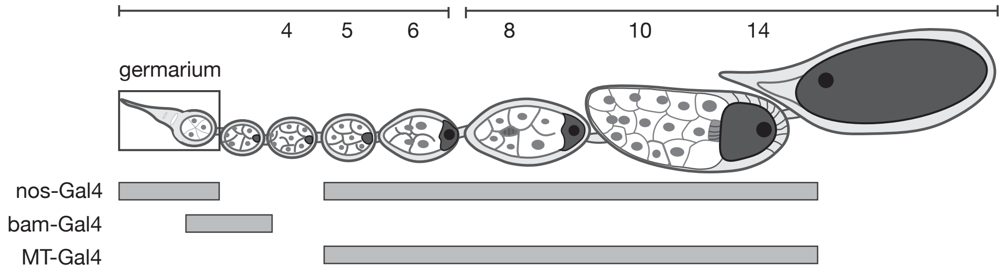


**B**

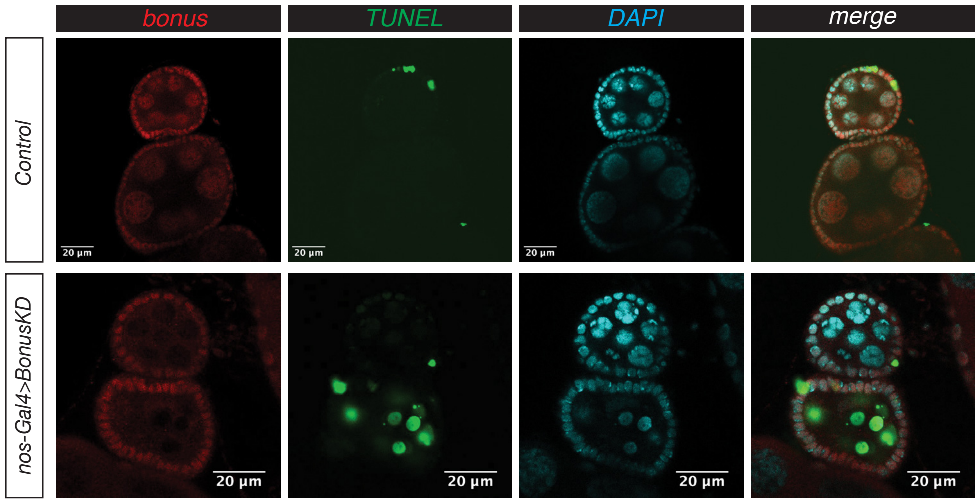


**C**

| Phenotype | Count |
|---|---|
| Normal | 8 (12%) |
| Hypomorphic | 16 (23%) |
| Rudimentary | 44 (65%) |

**D**

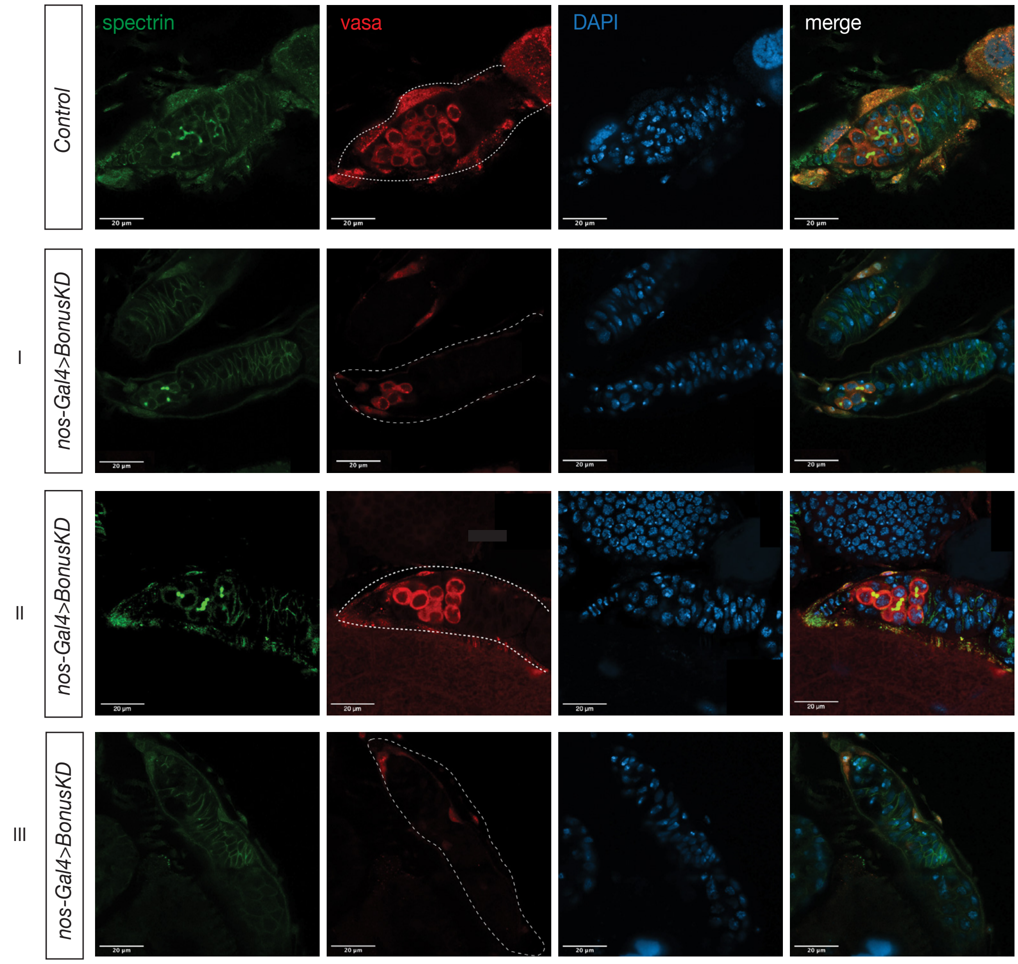


**E**

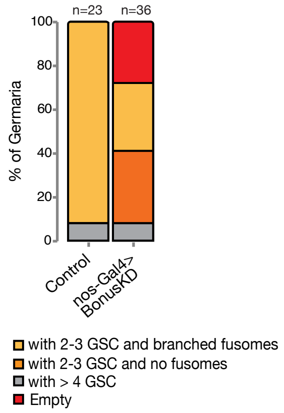

Extended Data Fig.2

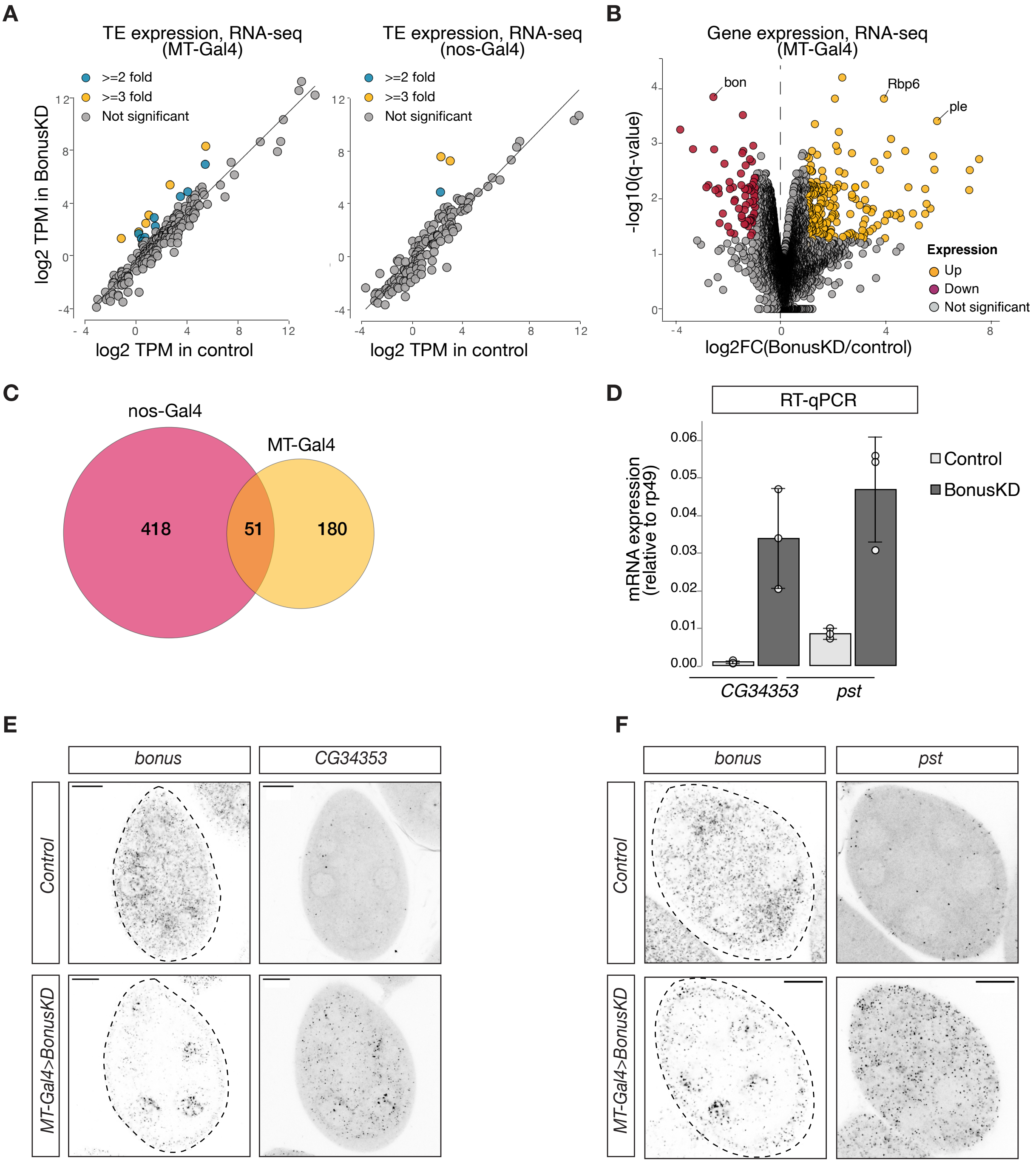

### Extended Data Fig.3

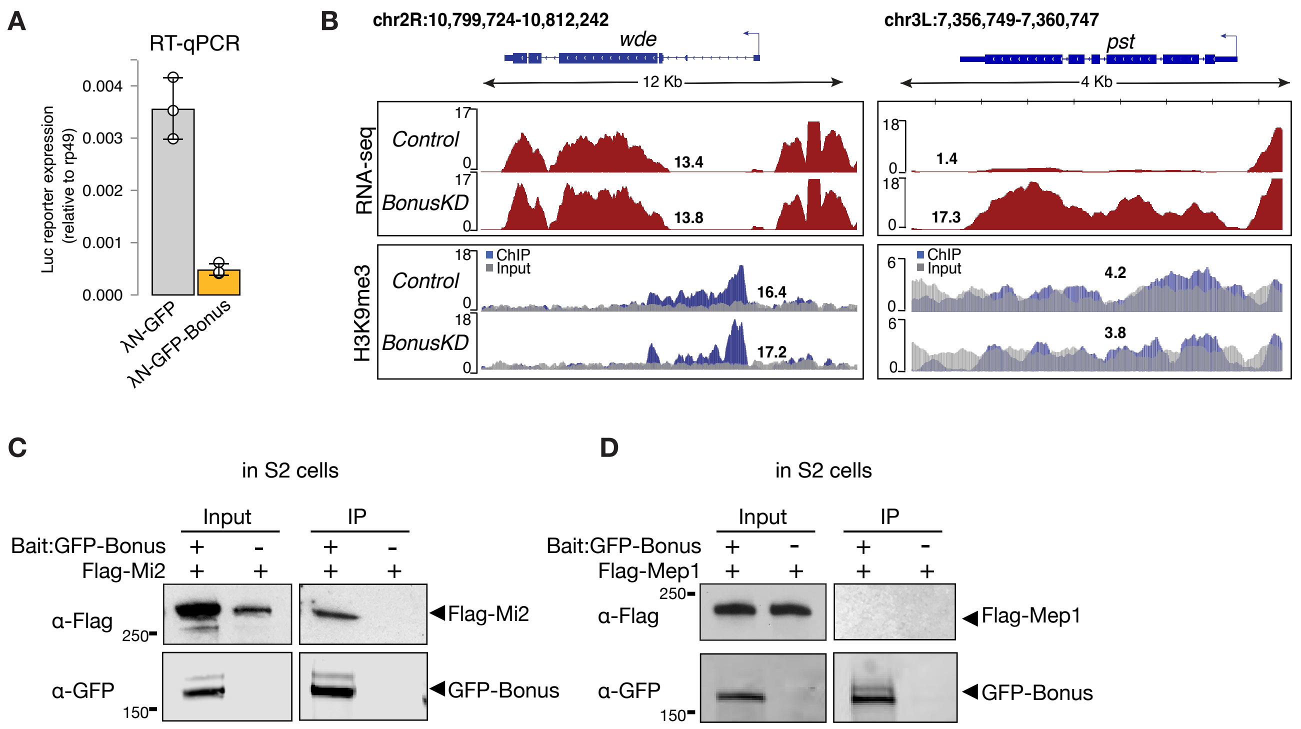

#### Extended Data Fig.4

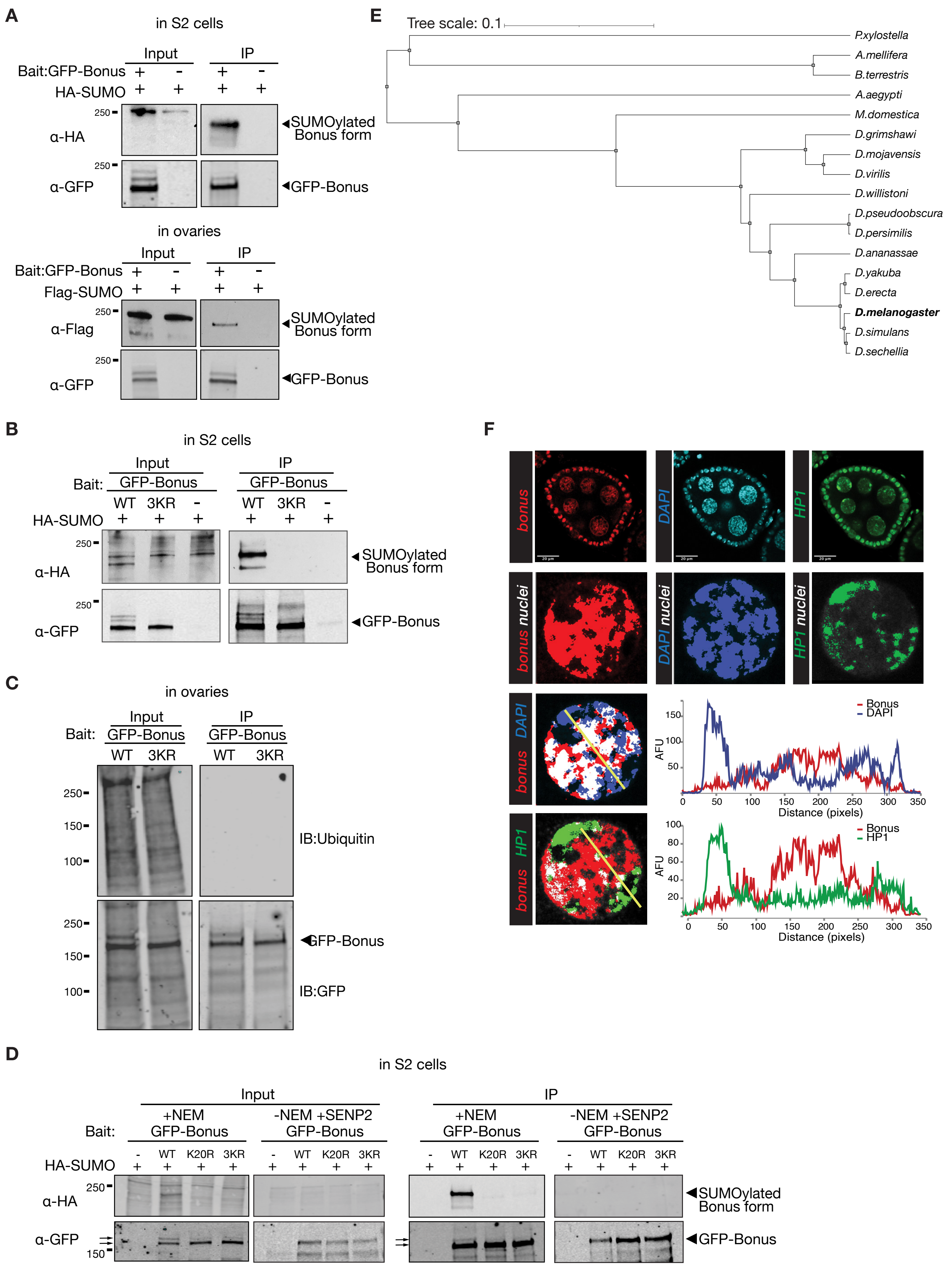

**Extended Data Fig.5**

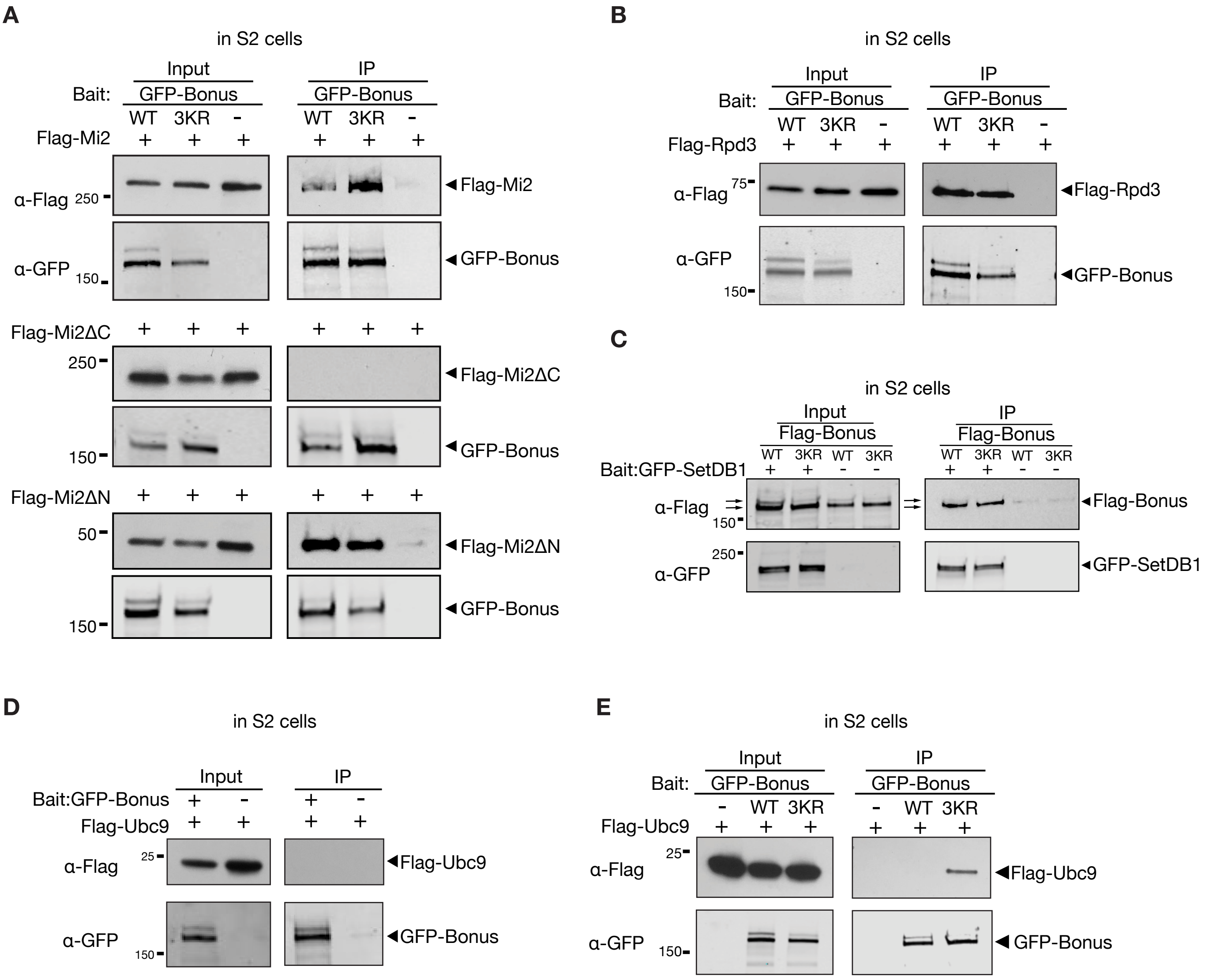
